# Virus-like DNA tracers for characterizing wastewater-based pathogen monitoring systems

**DOI:** 10.1101/2023.12.24.573197

**Authors:** Anjali Gopal, Ari N. Machtinger, Brian Wang, Daniel P. Rice, William J. Bradshaw, Michael R. McLaren, Kevin M. Esvelt

## Abstract

Wastewater-based pathogen monitoring has recently become a ubiquitous method for tracking the abundance and evolution of pathogens in a community or port of entry. But accurate inference of disease incidence from wastewater remains challenging, in large part due to noise and bias arising from wastewater transport and measurement. We developed a new system of tracer particles designed to enhance wastewater monitoring of viral pathogens. These tracers, which consist of synthetic DNA barcodes packaged into non-infectious viral capsids, can be deposited at known amounts and times into the sewer system and measured in wastewater samples to assess features such as assay sensitivity, and the decay and delay in viral signal arising from a shedding event. Here we describe the design, construction, and initial characterization of a 25-plex tracer library. Our tracer library supports simultaneous measurement of up to 5 distinct tracers by qPCR and up to 25 distinct tracers by amplicon or metagenomic sequencing. We demonstrate experimentally that the tracers are biologically inert and exhibit good stability and qPCR-assay performance in wastewater. These features collectively suggest that barcoded, virus-like tracers have the required properties for high-accuracy, high-resolution characterization of wastewater monitoring systems.

## Introduction

Rapid and accurate pathogen monitoring can improve public health responses to endemic and emerging diseases (Gardy and Loman 2018; Diamond et al. 2022; Hayman et al. 2023). Currently, most pathogen monitoring is based on either clinical diagnosis or detection of specific pathogens. These methods require individuals to seek care or participate in a testing program, limiting their reach (Islam et al. 2023). Public health coordinators must collect and aggregate the resulting data from many sources, which introduces further delays and complicates geographic comparisons.

A complementary approach is to monitor wastewater using molecular assays that detect DNA or RNA from pathogens shed into the sewer system by infected people (O’Keeffe 2021; Diamond et al. 2022). Wastewater monitoring has detected a wide range of pathogens, including influenza, polio, and norovirus (Toribio-Avedillo et al. 2023; Ryerson 2022; Lago et al. 2003; Boehm et al. 2023); these efforts increased dramatically during the COVID-19 pandemic to track the rate and genetic evolution of SARS-CoV-2 infections (Shah et al. 2022; Diamond et al. 2022). The choice of sampling point determines the breadth of the monitored population: sampling building outflows and manholes targets specific buildings or neighborhoods (Fahrenfeld et al. 2022); sampling at wastewater treatment plants (WWTPs) targets city subsections, including multi-city regions (Duvallet et al. 2022); and sampling from airports or aircraft targets the global travel network (Hjelmsø et al. 2019; Morfino et al. 2023). The choice of assay determines whether the monitoring is specific to predetermined pathogens (qPCR, amplicon sequencing) or able to detect a broad spectrum of pathogens (metagenomic sequencing). In this way, wastewater monitoring offers a non-invasive and potentially cost-effective method to track the abundance and genomic evolution of pathogens.

However, confidently predicting the level and distribution of infections in a given population using wastewater remains challenging. Quantitative precision is hampered by the tenuous link between measured pathogen concentrations in wastewater and community infections. The rate and duration of shedding into the sewer system can vary by individual and pathogen variant (Lavania et al. 2022; Prasek et al. 2023), but much of the uncertainty arises further downstream: Pathogens shed into the system at varying locations take different (and unknown) lengths of time to arrive at the sampling location, during which the detectable particles may decay and spatially disperse. Samples collected on different days may be impacted by varying environmental factors, such as dilution with stormwater and aggregation with non-human sewage. The molecular assays themselves may introduce further noise and bias, which may not be well-characterized by standard laboratory controls. There are currently no suitable control materials or techniques for characterizing and correcting the error in disease inference that arises from the aggregate effects of wastewater transport and measurement.

A candidate methodology to measure and reduce this error would involve depositing tracer particles at known amounts and times into different parts of the sewer system and monitoring their concentration in wastewater samples collected downstream (Figure 1). Tracers in the form of chemicals or synthetic DNA have previously been used to characterize natural and man-made water systems (Davis et al. 1980; Mikutis et al. 2018; Georgakakos, Richards, and Walter 2019; Pang et al. 2020). One study even conducted a limited characterization of a wastewater monitoring program for polio by flushing live attenuated poliovirus down a toilet (Hovi et al. 2001). Unfortunately, existing tracer systems either do not resemble real pathogens, are unable to be measured by standard molecular assays, cannot be distinguished from real infections, or don’t allow for multiple tracer types to be used in multiple locations.

**Figure 1.**
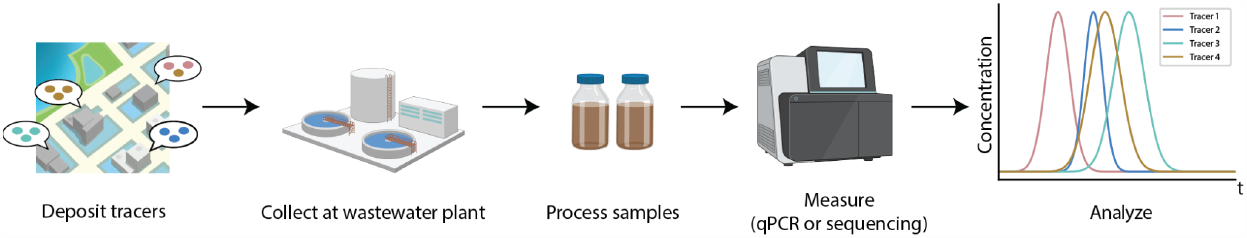
Overview of the tracer system. Tracers can be deposited into deposition sites (e.g., toilets) across many locations in a city. After deposition, wastewater samples can be collected from local treatment plants, and analyzed via qPCR or sequencing. By barcoding tracers, we can pool them together for detection in a single experiment.

To overcome these limitations, we developed a new form of tracer which consists of synthetic DNA packaged into non-infectious viral capsids. Like previous synthetic DNA tracers, our virus-like tracers contain unique DNA barcodes which can distinguish between depositions at different locations and different times. Unlike previous tracers, ours behave like real viral particles during wastewater transport, sample collection, and measurement. We achieve these improvements by packaging our tracers using the M13 bacteriophage, which we modified to contain DNA barcodes and to be biologically inert and environmentally harmless. Here we detail the design, construction, and initial characterization of these tracers.

## Results

### Design and construction of virus-like DNA tracers

We identified the following design requirements for an ideal tracer system to calibrate wastewater monitoring systems for viral pathogens: (1) Tracers should contain DNA barcodes that permit measurement via the three dominant measurement modalities—qPCR, amplicon sequencing, and metagenomic sequencing—for wastewater monitoring, with sufficient diversity between tracer “genotypes” to facilitate simultaneous deposition and measurement of multiple tracers across these measurement modalities. (2) Tracers should mimic naturally-occurring viruses during transit and measurement. (3) Tracers should be non-infectious and unable to replicate in the wastewater environment, yet easy to produce in the laboratory. Previous tracer systems could not satisfy all these requirements (Supplementary Note S1).

We based our tracers around a genetically engineered, selectively replicating phagemid packaged by the M13 bacteriophage (Figure 2A). M13 is a well-characterized phage that has been widely used for phage display applications; experiments have modified M13 to eliminate the need for helper phage by using helper plasmids instead (Chasteen et al. 2006). We leveraged these properties to develop tracer particles that only contain our “phagemid genome”, which include barcoded synthetic DNA. Furthermore, tracer particles were modified to be both non-infectious to M13’s host species, *Escherichia coli* (Figure 2B), and we additionally ensured that the phagemid genome would not be able to replicate itself within wild-type *E. coli* (Figure 2C).

**Figure 2.**
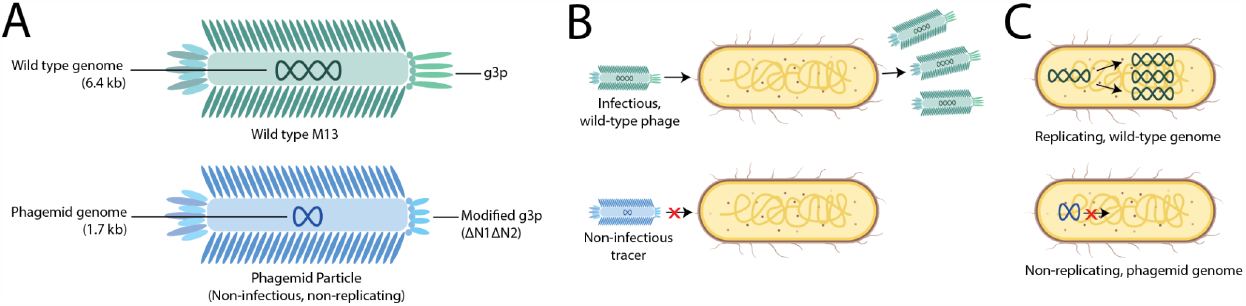
Overview of tracer particle design. A) Tracer particles are based on modified M13 phage particles (green: wild type, blue: phagemid), but with a truncated “phagemid genome”. B) The tracers are designed to be non-infectious: unlike wild-type M13 phage (top), the tracer particles are unable to infect host cells (*E. coli*, orange). C) The tracer genome is also non-replicating. Unlike the wild-type M13 genome (top), the tracer’s phagemid genome has been engineered to be unable to replicate within wild-type *E. coli* cells.

Each tracer genotype contains a distinct 300 bp synthetic DNA barcode, which supports measurement via qPCR and via amplicon sequencing (Figure 3A). The full barcode consists of a qPCR region (Figure 3B) and a sequencing region (Figure 3C). The qPCR region enables sensitive measurement of the concentration of individual tracer genotypes via TaqMan (probe-based) qPCR assays. The sequencing region enables simultaneous measurement of all tracer genotypes via amplicon sequencing, and of the total tracer concentration via a single TaqMan qPCR assay. All primer and probe sets were computationally designed to optimize on-target qPCR performance while minimizing the chance for off-target measurement of other biological sequences. Our final tracer library consists of 5 sets of distinct qPCR barcodes, and 25 sets of distinct sequencing barcodes. This library supports measurements of up to 5 distinct tracers in a single wastewater sample using qPCR and up to 25 distinct tracers using amplicon sequencing. Full details of the barcode design can be found in Materials and methods.

**Figure 3:**
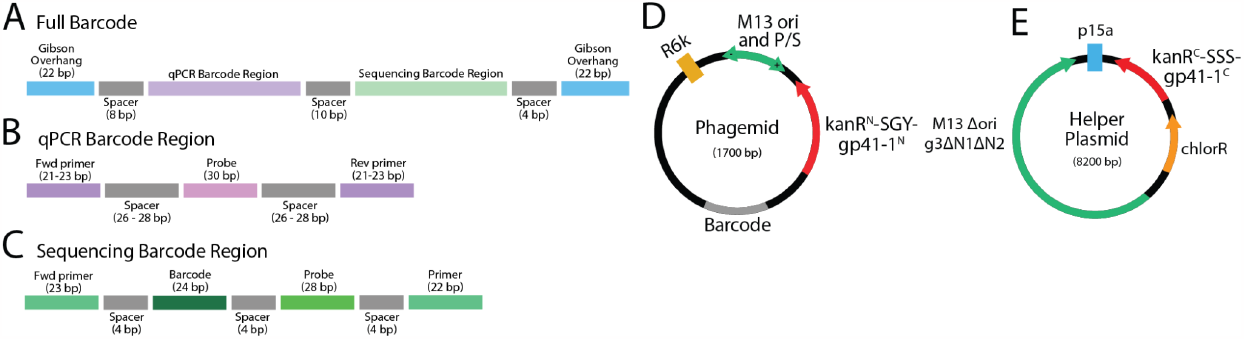
Barcode Design for Tracers. (A) Tracer barcodes consist of a qPCR barcode region and a sequencing barcode region, separated by spacers, and flanked by Gibson overhangs. (B) The qPCR barcode region consists of forward and reverse primer-binding sites and a probe-binding site, separated by spacers. Every unique qPCR barcode has unique primer-binding and probe-binding sites. (C) The sequencing barcode region consists of constant forward and reverse primer-binding sites, a unique barcode sequence, and a constant probe-binding site, separated by spacers. (D) The phagemid consists of the M13 origin (which also contains the packaging signal), an R6K origin, the N-terminal fragment of a split kanamycin resistance gene (kanR^N^-SGY-gp41-1^N^), and the synthetic barcode described in (A). (E) To generate non-infectious, non-replicating phagemid particles, we use a helper plasmid which contains the protein coding genes, with the N1 and N2 domains deleted from gene III, but does not contain the M13 origin or packaging signal. Instead, we use a p15A origin to replicate the plasmids in *E. coli*. The helper plasmid contains the C-terminal fragment of a split kanamycin resistance gene (kanR^C^-SSS-gp41-1^C^), as well as a chloramphenicol resistance marker for selection. Since the helper plasmid does not have the M13 origin or packaging signal, it is not packaged into the phagemid particle.

To render M13 tracer particles non-infectious, we create tracer particles using the helper plasmid system described in (Chasteen et al. 2006), wherein a phagemid plasmid contains the M13 origin of replication and packaging sequence, and the full tracer barcode (see Figure 3D). The remainder of the M13 phage genes are found in a helper plasmid (see Figure 3E). To render the tracer particles non-infectious, the N1 and N2 domains of protein III that contact the mating pilus and the TolQRA receptor used for entry (Kleinbeck and Kuhn 2021) were deleted from the helper plasmid backbone (Rakonjac, Jovanovic, and Model 1997). Furthermore, by utilizing the R6K origin of replication for the phagemid (see Figure 3D), the phagemid can only replicate within cells already containing the *pir* gene (Rakowski and Filutowicz 2013). Selection pressure to maintain both the phagemid and helper plasmid in the same cell line is provided by using the kanamycin-resistance intein system, wherein each plasmid contains one half of the intein needed to produce kanamycin resistance (Palanisamy et al. 2019). The use of split inteins avoids releasing full antibiotic resistance cassettes into the environment during deposition experiments.

### Non-replicability and non-infectivity

To assess the properties of our tracer system, we first performed assays to test for non-replicability and non-infectiousness. To test for replicability, we evaluated whether phagemids that were transformed into cell lines with and without the *pir* gene (pir-116 and DH10B cell lines, respectively) could replicate the plasmid and support kanamycin resistance. We hypothesized that cells without the *pir* gene would not support resistance. For this assay, we first transformed a quantification plasmid (Figure 4A) into the DH10B and pir-116 cell lines, which contains the C-terminal fragment of the kan intein (which is also found in the helper plasmid). The quantification plasmid-containing cell lines were then transformed with either a kanamycin-containing control plasmid (pHSG299) or the phagemid, and plated under kanamycin selection.

**Figure 4:**
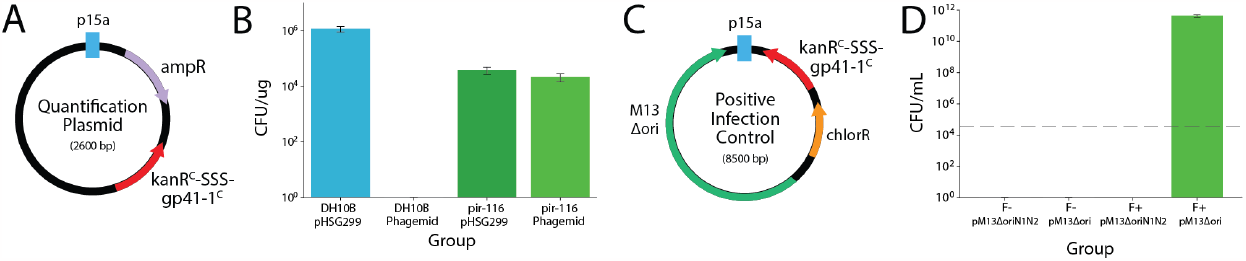
Assays assessing infectivity and replicability of phagemid particles. (A) A quantification plasmid is maintained in some cell lines for additional control and characterization experiments. The quantification plasmid contains the C-terminal fragment of a split kanamycin resistance gene (*kanR*^*C*^*-SSS-gp41-1*^*C*^), as well as an ampicillin resistance marker. (B) Non-replicability assays demonstrate that when the phagemid is transformed into either *pir-*expressing cell lines (pir-116) or control cell lines (DH10B), kanamycin resistance is only supported in the *pir*-expressing cell lines. Control experiments demonstrate that when kanamycin-containing control plasmid (pHSG299) is transformed into both pir-116 and DH10B cell lines, kanamycin resistance is supported in both cell lines. (C) To demonstrate non-infectiousness of our phagemid particles, we generate a strain of infectious particles using a positive infection control (PIC) plasmid to act as a positive control. The PIC plasmid has the N1 and N2 domains from gene III. (D) When infectivity assays are performed with pM13Δori particles (containing the N1 and N2 domains in gene III) or pM13ΔoriN1N2 particles (lacking N1 and N2), only F+ cell lines infected with pM13Δori particles support kanamycin resistance. Dashed gray line indicates limit of detection (< 3.3 × 10^4^ infectious particles per mL of total particles).

Our results indicate that the DH10B cell line supports kanamycin resistance when transformed with pHSG299 (∼1 × 10^6^ cfu/ug), but not when transformed with the phagemid (Figure 4B). The pir-116 cell lines support kanamycin resistance when transformed with both pHSG299 and the phagemid. These results support our hypothesis that the phagemid cannot be maintained in cell lines without the *pir* gene.

We next assessed the infectiousness of the tracer particles. For our infectivity assay, we grew two types of tracer particles: one which contained intact N1 and N2 domains in gene III (pM13Δori particles) grown using a “positive infection control” plasmid (PIC; Figure 4C), and one which contained N1- and N2-deleted gene III (pM13ΔoriN1N2 particles), grown using the helper plasmid. We then attempted infection of pir-116 cell lines either containing (F+) or lacking (F-) the F pilus, which is required to support phage infection. Our results indicate that only F+ cell lines infected with pM13Δori particles support kanamycin resistance (Figure 4D). Given an observed LLOD of < 3.3 × 10^4^ infectious particles per mL of total particles, and given an initial tracer particle titer of ∼10^12^ particles/mL, this suggests that fewer than 4 in 10^8^ of pM13ΔoriN1N2 tracer particles are infectious.

### Sensitivity and accuracy of tracer qPCR assay

To assess our ability to accurately detect the presence and quantify our tracers via qPCR, we conducted a spike-in experiment in which we varied tracer concentration in wastewater over many orders of magnitude. We spiked a single tracer genotype into 10 mL wastewater aliquots to produce concentrations ranging from approximately 90 particles/mL to 6×10^9^ particles/mL. We quantified the amount of tracer in each wastewater aliquot by concentrating and extracting DNA from the full 10 mL aliquot, then applying the tracer’s genotype-specific qPCR assay. We then analyzed the qPCR results to assess qPCR efficiency, linear dynamic range, and lower limit of detection of the assay.

Our results demonstrate high linearity of quantification within the range that we tested (6×10^9^ particles/mL to 90 particles/mL) with an R^2^ > 0.99% and high qPCR efficiency (Figure 5A). A subset of negative control samples, consisting of wastewater with no added tracer, showed amplification around cycle 38 (Supplementary Figure S1). Given that even our lowest concentration of spike in (90 particles/mL) had threshold cycle values below 38, we posit that the lower limit of detection of our assay is below 100 particles/mL.

**Figure 5:**
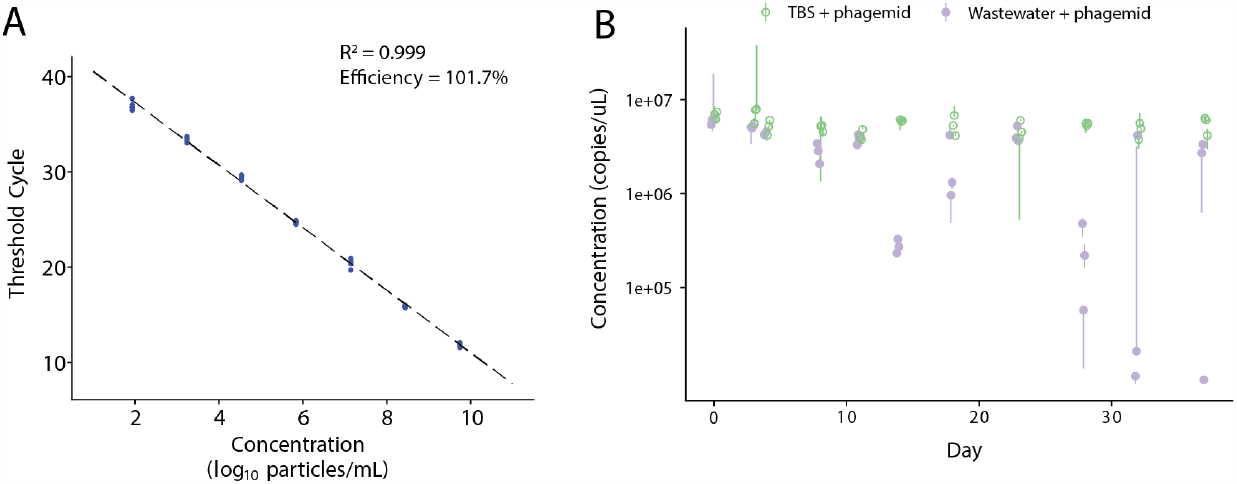
Assessing sensitivity and stability of phagemid particles. (A) When spiked into 10 mL of archived wastewater samples, phagemid particles demonstrated a linear dynamic range from 90 particles/mL to 6×10^9^ particles/mL when measured with qPCR. As a result, the LLOD of our assay is < 100 particles/mL. (B) A decay experiment measured the decay in tracer concentration (as measured by qPCR) over the span of 37 days. The figure shows each technical replicate with point and error bars representing the median and range of qPCR replicates, respectively. Tracers spiked into wastewater showed substantial decay between 10 to 37 days; however, high experimental variability limits our ability to make quantitative decay rate estimates. Tracers spiked into buffer showed negligible decay over the entire 37-day period.

### Stability of tracer particles

We next assessed the stability of the tracer particles in wastewater and in typical lab storage conditions. We spiked tracer particles into samples of archived wastewater and a standard lab buffer (TBS), stored the samples at room temperature, and quantified the tracer concentration via qPCR over a period of 37 days (Figure 5B). Tracers showed negligible decay in buffer over the entire 37-day period. In contrast, tracers appeared stable in wastewater only for a period of ∼10 days. Our results weakly suggest that tracers spiked into wastewater remain stable for ∼10 days, decline 10-fold by ∼20 days, and 100-fold by ∼30 days. However, high experimental variability limits our ability to draw strong conclusions about tracer particle decay in wastewater from this initial experiment.

## Discussion

Confident inference of community infection levels via wastewater monitoring requires appropriate experimental controls to assess noise and bias arising from wastewater transport and measurement. Tracer deposition experiments are promising for this purpose, but existing wastewater tracer particles are unsuitable. Here we developed a system of tracers, consisting of viral capsids containing a synthetic DNA barcode, to be used in multiplexed tracer deposition experiments. Each member of our 25-plex tracer library has a unique DNA barcode that enables simultaneous measurement of up to 5 distinct tracers by qPCR and 25 by amplicon or metagenomic sequencing. The tracers are biologically inert, environmentally harmless, and easy to produce and store in the lab. Together, these features make the tracers suitable for high-accuracy, high-resolution characterization of wastewater monitoring systems.

This tracer system offers a number of key advantages over previous tracer systems. In contrast to chemical tracers, synthetic DNA tracers like ours can be measured using the same types of molecular assays used to measure pathogens. Multiple tracer genotypes can be easily distinguished in a sample, enabling multiplexed deposition experiments. Furthermore, our tracers are also the first to support the simultaneous measurement of all tracers via amplicon sequencing in addition to measuring individual tracers via qPCR. Compared to previous synthetic DNA tracers, our tracers better resemble natural viruses and are easier to produce in the lab.

Our choice to base our tracers on the M13 virus in particular was driven primarily by the availability of appropriate genetic-engineering techniques. This choice, however, puts some limitations on the utility of the present tracer system. Our tracers, like the M13 bacteriophage they are based on, have a ssDNA genome and do not have a lipid envelope. Thus, they will not perfectly mimic pathogenic viruses that do not share these features. For instance, enveloped viruses such as SARS-CoV-2 may have different stability properties in wastewater, leading to different dynamics in wastewater transport and extraction. In addition, our tracers must be measured with qPCR and/or sequencing protocols suitable for ssDNA. They are not, for example, suitable for use as an internal control for metagenomic sequencing using the Illumina Nextera DNA Library Prep method, which only targets dsDNA. We believe our tracers can lay the groundwork for developing tracers that mimic other types of viruses.

Further work is needed to characterize and demonstrate the full utility of our tracers. For instance, we have yet to develop and validate an amplicon-sequencing protocol for simultaneous measurement of multiple tracers. In addition, our decay experiment showed high experimental variation in the wastewater treatment group, which limits our ability to estimate the decay rate of tracer concentration via qPCR. The current experiment was carried out in archived wastewater samples which were stored at 4°C for over 5 months, which we expect to be less biochemically active than fresh wastewater. We aim to repeat this experiment in fresh wastewater under better controlled conditions in order to more closely estimate the decay that would occur in an actual deposition experiment. Finally, additional optimization of protocols for producing and measuring tracers may be required for deposition experiments at the world’s largest treatment plants. A sufficiently high combination of total tracer amount deposited and sensitivity of the measurement assay are needed to ensure detection despite the fact that tracers will be diluted in samples collected at the treatment plant by a daily flow that can reach a billion liters per day.

We are currently in conversation with regulatory authorities to receive approval to conduct a tracer deposition experiment to characterize a municipal wastewater monitoring system. We aim to share the tracers with research and testing labs with the aim of improving viral wastewater monitoring world-wide.

## Materials and methods

See Supplementary Table S1 for all primer sequences, and Supplementary Table S2 for a complete list of reagents and equipment used in these experiments. All plasmid maps are given in Supplementary Table S3, full tracer sequences are in Supplementary Table S4, and tracer component sequences (including primer and probe sequences and 24-mer sequencing barcode) are in Supplementary Table S5.

### Barcode design

#### Overview

The full barcode consists of two regions, a qPCR region and a sequencing region, which are separated by spacer sequences and flanked by Gibson assembly overhangs (Figure 3). The qPCR region and the sequencing region each consists of an internal sequence flanked by a pair of forward and reverse primer sequences. The internal sequence contains a probe sequence for use with TaqMan qPCR measurement and, for the sequencing region only, an additional 24-mer sequencing barcode unique to each tracer genotype. Since there are 5 unique qPCR barcode sequences in our library, each with a unique primer-probe set, we can simultaneously measure up to 5 different tracers in a pooled sample using TaqMan qPCR. Similarly, each phagemid has a unique sequencing barcode, which enables measurement of all 25 tracers simultaneously with a combination of amplicon sequencing and a single qPCR reaction. Each sequencing-barcode sequence has the same qPCR primer-probe set, but a unique 24-mer internal barcode; thus qPCR can be used to measure the total concentration of all tracers, whereas amplicon sequencing can be used to determine the relative abundances of the 25 genotypes based on their distinct 24-mer barcodes.

We randomly generated primer and probe pairs using a modified version of the barCoder software package (Bernhards et al. 2021). We modified barCoder to skip the built-in BLAST step, in order to more quickly generate candidate sequences. As a replacement, we later used NCBI’s Primer-BLAST (Ye et al. 2012) to computationally verify that our primers were not predicted to amplify known sequences.

Sequencing barcodes are 24-mer sequences corresponding to one of the distinct Oxford Nanopore Technologies (ONT) barcodes described in the adapters section of the Porechop custom library (https://github.com/rrwick/Porechop/; Wick et al. 2017). We used these barcodes due to their tolerance for high sequencing error rates, enabling accurate barcode assignment even with the higher-error sequencing platforms such as the ONT Minion.

Final qPCR and sequencing regions were generated from barCoder output with custom post-processing to adjust spacers and, for the sequencing region, to insert the 24-mer internal barcode; see code on [GitHub](https://github.com/naobservatory/phagemid-tracers/tree/main/phagemid-design/analysis). For the qPCR region, primer (21 - 23 bp) and probe (30 bp) sequences are separated by spacer regions of 26 - 28 nucleotides. The sequencing barcode region has a varying internal barcode sequence (24 bp) and a constant probe sequence (28 bp), surrounded by constant primer sequences (22 - 23 bp) separated by 4 - 5 bp spacer sequences.

We determined the final barcode sequences using an interactive design and laboratory-evaluation process. First, we tested 20 candidate primer-probe sets, picking 5 to serve as the qPCR barcode sequences and 1 to provide the primer-probe set for the sequencing barcode. We then generated 96 phagemid strains, sequenced their barcode regions, and picked 25 strains so as to have 25 distinct barcode sequences, with 5 sequencing barcodes per qPCR barcode (see Library Construction and Validation). The full sequences for the final 25 tracers are in Supplementary Table S4 (Full Tracer Sequences) and Supplementary Table S5 (Barcode Sequences and Components). The naming convention for barcodes is: “T1 (Tracer Tag 1) _ QS2 (qPCR barcodes from set 2) _ XX (qPCR barcode ID) _ BCYY (porechop barcode ID)”.

### Plasmid construction and bacterial cloning

#### Helper plasmid and positive infection control

M13cp plasmid in a DH5αF′ bacterial host was constructed in-house based on previously published sequences (Chasteen et al. 2006). pSiMPlk was obtained from Addgene (Palanisamy et al. 2019). The M13 phage genome was amplified in two parts: part 1 used primers M13cp_part1_fwd and M13cp_part1_rev, and part 2 used primers M13cp_part2_fwd and M13cp_part2_rev. kanR^C^-SSS-gp41-1^C^ was amplified from pSiMPlk_C using forward and reverse primers KanR_gp41-1c_part1_fwd and KanR_gp41-1c_part2_rev. The complete list of primer sequences used in this study is given in Supplementary Table S1, and all plasmid maps can be seen in Supplementary Table S3. After Gibson assembly was complete, a second PCR was performed using primers HPsig_fwd and HPsig_rev, and a final one-piece Gibson assembly was performed.

For the positive infection control, one linear PCR was performed on M13cp with forward and reverse primers M13cp_part1_fwd and M13cp_part2_rev, and Gibson assembly was performed with the same kanR^C^-SSS-gp41-1^C^ gene, using the methods described above.

#### Quantification plasmid

The p15A origin was amplified from M13cp using forward and reverse primers M13cp_qp_fwd and M13cp_part2_rev. The ampicillin resistance gene was amplified from a custom in-house plasmid using forward and reverse primers AmpR_qp_fwd and AmpR_qp_rev. kanR^C^-SSS-gp41-1^C^ was amplified from pSiMPlk using forward and reverse primers kanR_qp_fwd and KanR_gp41-1c_part2_rev.

#### Phagemid

The R6k origin was amplified from a custom in-house plasmid and sequence verified, using forward and reverse primers R6k_fwd and R6k_rev. The f1 origin was amplified from M13cp using forward and reverse primers M13ori_fwd and M13ori_rev. kanR^N^-SGY-gp41-1^N^ was amplified from pSiMPlk_N using forward and reverse primers kanR_gp41-1N_fwd and kanR_gp41-1N_rev. For barcodes, custom sequences were ordered from IDT and designed as specified in the “Barcode Design” section above.

All fragments were PCR amplified using either PrimeStar Max (for fragments < 2 kb) or PrimeStar GXL (for fragments > 2kb), purified via gel extraction, and digested via Dpn1. Plasmids were assembled via Gibson assembly using the NEBuilder HiFi DNA Assembly Master Mix; parts were pooled in equimolar concentrations of up to 0.2 pmol per fragment, and incubated in 20 uL at 50°C for 15 minutes.

All DNA cloning was performed using pir-116 cells or DH10β cells. For electroporation, the Eppendorf Eporator Electroporator was used with a 2000V charge. 50 - 100 μL of cells were transformed with 1 μL of plasmid (either straight from the Gibson cloning mixture, or from mini-prepped DNA). Cells were then immediately incubated in S.O.C. medium for 1 hour in an incubator at 37°C and 230 rpm.

For helper plasmid or PIC-containing cell lines, pir-116 cells were transformed via electroporation with the assembled helper plasmid or positive infection control, and then plated on LB agar plates containing 40 mg/mL of chloramphenicol. For quantification plasmid-containing cell lines, pir-116 cells were transformed via electroporation with the assembled quantification plasmid and then plated on LB-agar plates with 50 mg/mL of carbenicillin. Single colonies were used to inoculate fresh cultures, and sequence verified via Nanopore sequencing.

### Library construction and validation

To prepare phagemid libraries, initial confirmation of the phagemid design was performed by cloning the phagemid backbone (containing all components except for the tracer barcode) into a helper plasmid containing cell line. The phagemid was transformed via electroporation as described above, and after a 1-hour recovery period, plated on LB-agar plates with 15 ug/mL of kanamycin. Single colonies were used to inoculate fresh cultures, and sequence verified via Nanopore sequencing.

To create the phagemid library, the phagemid backbone was linearized using PCR via forward and reverse primers tag_overhang_fwd and tag_overhang_rev. Custom barcodes were ordered from IDT and designed as specified in the “Barcode Design” section above. 96 distinct eBlocks, consisting of barcodes with 5 unique qPCR regions distributed across 96 unique sequencing barcodes were ordered from IDT and designed as specified above. 1 μL from each eBlock was pooled into a single mastermix and used in a Gibson assembly method, as described above. The assembled phagemid DNA was then transformed into a quantification plasmid-containing cell line via electroporation and plated on LB-agar plates with 15 ug/mL of kanamycin. After overnight growth, 96 single colonies were used to inoculate fresh cultures, and were then sequence verified. From the sequence verified results, 25 unique strains were picked, to yield a library with tracers that had 5 unique qPCR barcodes, and 25 unique sequencing barcodes.

The 25 unique phagemid-containing quantification plasmid cell lines were mini-prepped and subject to restriction digestion via *PstI* and *ScaI*. The remaining DNA from each strain was transformed into the helper plasmid-containing cell lines via chemical transformation. After overnight growth, single colonies were used to inoculate fresh cultures, and used to create the final library. In addition, a single infectious phagemid particle strain was also made by transforming the TS1_QS2_010_BC33 phagemid into a PIC-containing cell line.

See Supplementary Table S1 for all primer sequences, and Supplementary Table S2 for a complete list of reagents and equipment used in these experiments. All plasmid maps can be seen in Supplementary Table S3, and full tracer sequences can be in Supplementary Table S4.

### Tracer particle isolation and purification

Each specific strain of tracer was grown in an individual bacterial culture in 2 - 5 mL of LB media with kanamycin (3015 ug/mL). After overnight incubation at 37°C and 250 - 300rpm, bacterial cultures were pelleted by centrifugation at 13,000 x g for 2 minutes. 1.2 mL of the supernatant was transferred to a fresh microcentrifuge tube and incubated with 300 μL of 20% PEG-8000, 2.5M NaCl solution. Viral particles were then pelleted by centrifugation at 13,000 x g for 3 minutes. Supernatant was discarded, and viral particles were pelleted again at 13,000 x g for 1 minute; residual supernatant is discarded with a 100 μL tip. The pellet was resuspended in 120 μL of 1X TBS by vigorous vortexing and incubated on ice for 1 hour. After incubation, the solution was vortexed again and centrifuged at 13,000 x g for 1 minute. The supernatant, which contains the purified viral particles, was transferred to a fresh tube. Viral particle concentration was measured via qPCR using the TaqPath ProAmp Mastermix, using a standard curve made from ssDNA that was quantified using the QuantIt ssDNA assay.

### Tracer particle DNA extraction

Cell suspensions or wastewater samples containing tracer particles were centrifuged at 5000 x g for 15 minutes at room temperature. Supernatants containing the tracer particles were transferred to a fresh microcentrifuge tube. The centrifugation step was repeated to reduce bacterial cell carryover. A 1:100 volume of Buffer MP (3.3 of citric acid monohydrate in 3 mL of water, dissolved and filtered through a 0.2 um sterile filter) was added to the supernatant. The solution was loaded onto a QIAprep spin column, and either vacuum filtered or centrifuged at 8000 rpm for 15 s. Flow-through was discarded. 0.7 mL of Buffer PB from the QIAprep Spin Miniprep Kit was added to the spin filter to enable M13 lysis and binding, and either vacuum filtered or centrifuged as described previously. Another 0.7 mL of Buffer PB was added, and the sample was incubated for 1 min to facilitate lysis, and then vacuum filtered or centrifuged described above. The spin column was then washed with 0.7 mL of buffer PE from the QIAprep Spin Miniprep Kit. Finally, 100 μL of elution buffer (either 10 mM Tris-HCl, pH 8.5 or nuclease-free water) was loaded into the middle of the spin filter, and the sample was incubated for 10 minutes at room temperature. After incubation, the spin column was loaded onto a fresh microcentrifuge tube and centrifuged at 8000 rpm for 30 s to elute the tracer DNA.

### Replicability assays

Pir-116 (Thermo Fisher Scientific) and DH10B (Thermo Fisher Scientific) cell lines were transformed with quantification plasmid via electroporation, as described above. Overnight cultures of single colonies grown in 2XYT were supplemented with kanamycin were grown at 37 °C with shaking at 230 r.p.m. to an optical density (OD600) of roughly 0.6–0.8. Cells were made electrocompetent via serial washing in 10% glycerol. 50 μL of electrocompetent cells were transformed with 100 ng of DNA via electroporation at 2000V with either a kanamycin-containing control plasmid (pHSG299) or phagemid, respectively. Cells were placed in 1 mL of S.O.C. medium with a 1 hour recovery time. 100 μL volumes of 1:10 and 1:100 dilutions of transformants were plated on LB-Agar plates with kanamycin. Plates were incubated at 37°C for 12-16h and checked for colony-forming units.

### Infectivity assays

Pir-116 cell lines were first conjugated with the F-plasmid via the S2060 strain expressing RFP. After conjugation, cells were plated on LB-agar with carbenicillin and tetracycline, and non-RFP expressing colonies were used to inoculate fresh cultures. F’ pir-116 cell lines were made electrocompetent via successive washes in 10% glycerol, and were transformed with quantification plasmid.

Non-infectious phagemid particles from the TS1_QS2_010_BC33 strain (pM13ΔoriN1N2 particles) were purified as described above. Infectious particles (pM13Δori particles) from the TS1_QS2_010_BC33 strain cultured from the positive infection control cell line were also purified as described above.

10 sets of 1:10 serial dilutions of both pM13ΔoriN1N2 and pM13Δori particles were initially performed. An initial infectivity assay was then performed using both the F+ and F-quantification plasmid cell line by spiking in 20 uL of the tracer particles to 180 uL of cell suspension. Spiked-in cell lines were incubated for 1 hour at 37°C and 230 rpm, and plated in 3 uL spots on LB-agar plates with kanamycin (30 ug/μL). Plates were incubated at 37°C for 12-16h and checked for colony-forming units. Following this initial assay, both pM13Δori and pM13ΔoriN1N2 particles were further diluted to 6-log_10_ and 7-log_10_ dilutions, and spiked into both the F+ and F-quantification plasmid cell line at a 1:10 ratio, and then plated on LB-agar plates with kanamycin (30 ug/μL) at 100 uL volumes. Plates were incubated at 37°C for 12-16h and checked for colony-forming units.

### Stability assay

Tracer particles from the T1_QS2_018_BC04 strain were purified as described above. Tracer particle concentration was determined via qPCR (data not shown) and tracers were diluted to a starting concentration of 1 × 10^6^ particles per uL. Archived wastewater samples were obtained from a local treatment plant, and stored at 4°C for five months. 60 mL of wastewater was withdrawn from an initial 500 mL aliquot and spiked with 15.9 μL of tracer particles. The spiked-in wastewater aliquot was then split into several 700 μL samples. Another 60 mL wastewater aliquot, without tracer particle spike-in, was used as a negative control to assess background signal during every time point. A TBS aliquot, with tracer particle spike-in, was used as another negative control to assess background rate of tracer degradation in buffer.

Timepoints for sampling are listed in Supplementary Table S4 and ranged from once per day to once per week over the course of the experiment. At each time point, DNA was extracted from a single sample from each of the three treatment conditions as described in Tracer particle isolation and purification and stored at -80C until the end of the experiment. Afterwards, qPCR was performed on all extracted samples.

### Spike-in experiment

Tracer particles from the T1_QS2_010_BC33 strain were purified as described above and concentration was determined via qPCR. Tracers were diluted to a starting concentration of 6.4 × 10^8^ particles/μL and diluted in serial 1:20 dilutions down to 0.1 particles/uL, for a total of 7 dilutions (see Supplementary Table S6 for the full list of dilutions and concentrations). Archived wastewater samples, obtained from a local treatment plant seven months ago, were stored at 4°C. 24 × 10 mL aliquots were withdrawn from the same 500-mL wastewater aliquot. For each tracer particle dilution, 3 replicates of 100-μL spike-ins were performed. Triplicate negative controls, consisting of wastewater with no phagemid spike in, were also set aside. After spike in, all wastewater samples were vortexed for 30s to ensure adequate mixing of tracer particles in wastewater.

All wastewater samples were vacuum filtered through a 0.22 um PES membrane. After filtration, samples were concentrated via Amicon Ultrafiltration with a 30 kDa membrane at 1500 g for 30 minutes. After concentration, the retentate was extracted, and phagemid DNA was purified as described in the “Tracer particle DNA extraction” section. The concentration of T1_QS2_010_BC33 was measured via qPCR.

## Supporting information

Supplementary Table S1: Primer Sequences

Supplementary Table S2: Complete list of reagents, equipment, and suppliers

Supplementary Table S3: Plasmid Maps

Supplementary Table S4: Full Tracer Sequences

Supplementary Table S5: Barcode Sequences and Components

Supplementary Table S6: Dilutions and concentrations for spike-in experiment

## Data and code availability

Code and output for designing the tracer barcodes and for analyzing the stability experiment is available in a GitHub repository (https://github.com/naobservatory/phagemid-tracers) and Zenodo archive (https://doi.org/10.5281/zenodo.10427946).

## Author contributions

K.M.E, M.R.M., A.G., and W.J.B. conceived the study; K.M.E., A.G., and M.R.M. designed the tracers with feedback from B.W.; A.G., B.W., and A.M. performed all experiments with feedback from K.M.E.; M.R.M., A.G., and D.P.R. performed all analysis; and A.G. and M.R.M. drafted the manuscript, with feedback from all authors. All authors have approved the final version of the manuscript.

## Acknowledgements

B.W. was supported by an Early-Career Funding for Global Catastrophic Biological Risks scholarship from the Open Philanthropy Project.

We thank Laura Sowin, Summer DeAmelio, and Boqiang Tu for helpful discussions with experiments and assistance with sequencing validation. We thank Amy Xiao for helpful discussions around wastewater processing methodology.

Schematics and vector art in Figures 1 and 2 were created with DALL·E 3 and Biorender.com.

## Supplementary Information

### Supplementary tables

Supplementary tables can be found in the supplementary tables folder in the manuscript’s GitHub repository ([link](https://github.com/naobservatory/phagemid-tracers/tree/main/manuscript/supplementary-tables)) and the associated Zenodo archive ([link](https://github.com/naobservatory/phagemid-tracers/tree/main/manuscript/supplementary-tables)). All tables are provided as TSV files which contain all essential information; in some cases, an Excel file is also provided to additional preserve formatting or equations.

Table S1: Primer Sequences :: [TSV](https://github.com/naobservatory/phagemid-tracers/blob/main/manuscript/supplementary-tables/Supplementary%20Table%20S1%20-%20Primer%20Sequences.tsv)

Table S2: Complete list of reagents, equipment, and suppliers :: [Excel](https://github.com/naobservatory/phagemid-tracers/blob/main/manuscript/supplementary-tables/Supplementary%20Table%20S2%20-%20Reagents%20Equipment%20and%20Suppliers.xlsx), [TSV](https://github.com/naobservatory/phagemid-tracers/blob/main/manuscript/supplementary-tables/Supplementary%20Table%20S1%20-%20Primer%20Sequences.tsv)

Table S3: Plasmid Maps :: [TSV](https://github.com/naobservatory/phagemid-tracers/blob/main/manuscript/supplementary-tables/Supplementary%20Table%20S1%20-%20Primer%20Sequences.tsv)

Table S4: Full Tracer Sequences :: [TSV](https://github.com/naobservatory/phagemid-tracers/blob/main/manuscript/supplementary-tables/Supplementary%20Table%20S1%20-%20Primer%20Sequences.tsv)

Table S5: Barcode Sequences and Components :: Excel, [TSV](https://github.com/naobservatory/phagemid-tracers/blob/main/manuscript/supplementary-tables/Supplementary%20Table%20S1%20-%20Primer%20Sequences.tsv)

Table S6: Dilutions and concentrations for spike-in experiment :: [Excel](https://github.com/naobservatory/phagemid-tracers/blob/main/manuscript/supplementary-tables/Supplementary%20Table%20S2%20-%20Reagents%20Equipment%20and%20Suppliers.xlsx), [TSV](https://github.com/naobservatory/phagemid-tracers/blob/main/manuscript/supplementary-tables/Supplementary%20Table%20S1%20-%20Primer%20Sequences.tsv)

### Supplementary Note S1: Previous tracer systems

Tracers are a well-established method of characterizing the movement and loss of particles in a water system (Davis et al. 1980). Chemical tracers (dyes and salts) and synthetic DNA tracers have been used to characterize natural water systems (Mikutis et al. 2018; Pang et al. 2020) and to track septic pollution (Georgakakos, Richards, and Walter 2019). Biological tracers in the form of naturally occurring microorganisms and viruses have also been used (Davis et al. 1980), including the use of live attenuated poliovirus by Hovi et al. 2001 to semi-quantitatively characterize polio monitoring at a wastewater treatment plant.

However, existing tracer systems are limited in many ways. First, while synthetic DNA and biological tracers can both be measured with techniques that are also used to detect pathogens from wastewater—such as qPCR and sequencing—raw synthetic DNA tracers are degraded much more rapidly than packaged viruses (Pang et al. 2020). Encapsulated tracers, such as DNA encapsulated in silica or alginate-chitosan, provide additional stability; however, accessing the DNA within the tracers require dissolution by harsh chemical reagents, which can confound the tracer’s utility in mimicking the physicochemical properties of pathogens (Mikutis et al. 2018; Pang et al. 2020). Inactivated or attenuated natural pathogens cannot be distinguished from viruses shed by infected individuals and lack sufficient multiplexing capabilities (enabling multiple tracers to be distinguishable when deposited at multiple entry points in the same wastewater system). Furthermore, the possibility of incomplete attenuation or inactivation risks the undesired spreading of infectious particles through the wastewater system.

### Supplementary figures

**Figure S1:**
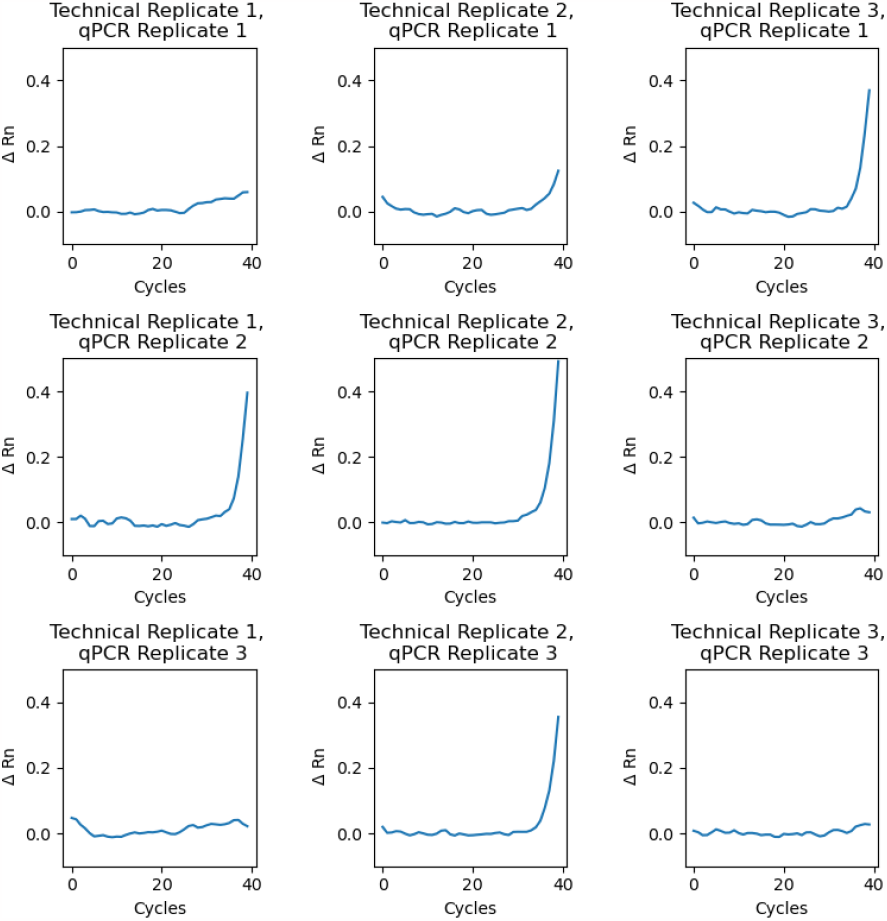
Amplification of Negative Controls during qPCR. Negative controls from spike-in experiments (containing wastewater but no phagemid particles) demonstrate some amplification. Samples consist of three technical replicates (columns). Each technical replicate has three independent qPCR replicates (rows). Even across technical replicates, not all qPCR replicates demonstrate amplification.

